# Assessing the Performance of LLMs in Multimodal Information Extraction for Biological Research: A Case Study on LLPS

**DOI:** 10.1101/2025.10.29.685285

**Authors:** Ka Yin Chin, Satoru Fujii, Shoichi Ishida, Kei Terayama

## Abstract

Advances in experimental techniques have expanded the volume of biological data. This has increased the demand for structured information extraction from papers, with large language models (LLMs) considered promising. However, challenges remain, including limited validation in biology and unclear applicability to multimodal tasks that integrate text with domain-specific figures, such as microscopic images and scatter plots. Here, we developed a multimodal LLM (MLLM)-based workflow to extract the experimental conditions and phase status from the text and figures of experimental papers on liquid-liquid phase separation and validated the effect of various inputs, prompts, and MLLM types. As a result, the Gemini 2.5 Pro-based extraction achieved an F1-score of 0.847 by processing each figure as a processing unit and inputting domain-specific prompts reflecting manual extraction guidance. This study demonstrates the potential and limitations of MLLMs for extracting biological information and provides insights for advancing multimodal approaches in biology.

## 1. Introduction

Advancements in experimental techniques in biology have led to a marked increase in data, promoting data-driven research. The biological literature indexed in PubMed continues to grow steadily and currently exceeds 38 million publications^1^. To make such knowledge accessible and usable, efforts have been made to structure information from literature and record it in databases^2–10^. For example, the Protein Data Bank (PDB) records three-dimensional protein structures^2^, GenBank aggregates gene and transcript information^3^, JASPAR provides transcription factor binding sites^4^, and BioGRID contains experimentally validated protein–protein interactions^5^. In addition, curation-based databases have also been maintained through expert annotation, such as KEGG^6,7^, ChEMBL^8,9^, and UniProt^10^, which rely heavily on manual extraction and annotation of unstructured knowledge from literature and external sources. The accumulated knowledge in such databases has been fundamental to recent advances in data-driven biology, enabling breakthroughs such as AlphaFold for protein structure prediction^11^, network-based approaches for functional inference^12^, and multi-omics integration for disease biology^13^. However, the success of these data-driven approaches relies on the knowledge structured in the databases, and manual curation remains a time-consuming and costly process that poses a bottleneck.

In recent years, information extraction from literature using large language models (LLMs) has been gaining attention to address these issues. Various domain-specific extraction approaches have been investigated, particularly in materials science and clinical medicine, and their utility has been consistently reported. In materials science, models such as MatSci-NLP^14^, MaterialsBERT^15^, and ChemDataExtractor^16^ have been developed to automatically extract material information such as synthesis conditions, crystal structures, and property values. These approaches have been applied to semi-automated knowledge base construction, further supporting the discovery of new materials and optimization of experimental conditions. In clinical medicine, methods have been developed to efficiently extract information about diseases, drugs, genes, and histopathological findings from biomedical documents^17–19^.

However, two major challenges prevent the application of these tools in the biological sciences. First, their development has primarily focused on materials science and clinical medicine with relatively few applications in biology. In materials science, databases remain relatively inadequate compared to the rapidly increasing number of papers, creating a strong motivation for developing automated extraction tools to advance data-driven research^20^. In clinical medicine, disease information is accumulated in natural language within electronic health records and similar systems, increasing the need for automated extraction technology to streamline its analysis and integration^21^. In contrast, for many classical topics in biology, the practice of recording information in existing databases has become routine, resulting in a relatively low immediate demand for extracting information from the voluminous literature. Nevertheless, the situation is now changing in emerging fields such as liquid-liquid phase separation (LLPS), single-cell biology, and microbiome research^22–24^. For example, in the case of LLPS, which is a highly important phenomenon where proteins and RNAs form condensates in response to environmental changes, the number of research papers has been increasing annually because of its relevance to diverse biological processes and neurodegenerative diseases^25–27^. This has led to the emergence of manually curated databases, such as LLPSDB^28^, RNAPhaSep^29^, and RNAPSEC^30^, which have enabled multiple data-driven studies^30–33^. However, the process of storing experimental data has not become routine, and updates require manual review by third-party research groups. Consequently, keeping pace with the growing volume of research output is difficult.

Second, the applicability of these LLMs to multimodal information extraction tasks, which require the integration of visual data such as microscope images and scatter plots with textual information, is yet to be thoroughly validated. To correctly extract information from biological literature, both visual elements and their surrounding textual context must be comprehensively understood. For example, in scatter plots, interpreting each data point requires correlating the numerical scales on the vertical and horizontal axes with the marker shapes in the legend and integrating the information from the caption and methodology section. In microscopic images, findings must be correctly understood by combining visual observations of the subject within the image with textual information. Although several attempts at data extraction have been made in the biological sciences in recent years^34, 35^, detailed performance evaluations of multimodal data extraction, including microscope images and scatter plots, have not been sufficiently conducted.

To overcome these challenges, this study aims to develop and evaluate information extraction methods from multimodal sources in the biological literature using LLPS as a case study. LLPS was selected because it is an important biological phenomenon, and environmental factors affecting the regulatory mechanism are recorded as experimental information in the text and figures, such as scatter plots and microscope images. In this study, we developed an extraction workflow for experimental conditions and phase information in each experiment using multimodal LLMs (MLLMs) and systematically examined the effects of various factors on the performance of extracting information related to scatter plots and microscope images. Specifically, the extraction processing unit, input data format, prompt design, and MLLM family were investigated. The results showed that a relatively high extraction performance was achieved using Gemini 2.5 Pro as the foundation, incorporating domain knowledge (DK) and extraction guidance into the prompt, feeding each figure with its associated text, and treating the figure as the processing unit. Furthermore, comprehensive analyses of positive and negative examples clarified the strengths of this MLLM-based approach and areas for future improvement.

## 2. Results

### 2.1. Overview of the extraction method

In this study, we developed an MLLM-based workflow to extract six experimental information, including protein name, protein concentration, RNA concentration, temperature, buffer pH, and phase state, from text and figures in experimental papers on LLPS. As shown in Figure 1, the overall workflow comprises three main steps: (1) preparation of multimodal input data, (2) information extraction using MLLMs, and (3) evaluation of the extracted results. The details for steps (1)-(3) are described in the method section. The extraction methods were evaluated using experimental information derived from 20 papers^36–55^ registered in RNAPSEC^30^. The details of the evaluation dataset are also mentioned in the method section.

**Figure 1.**
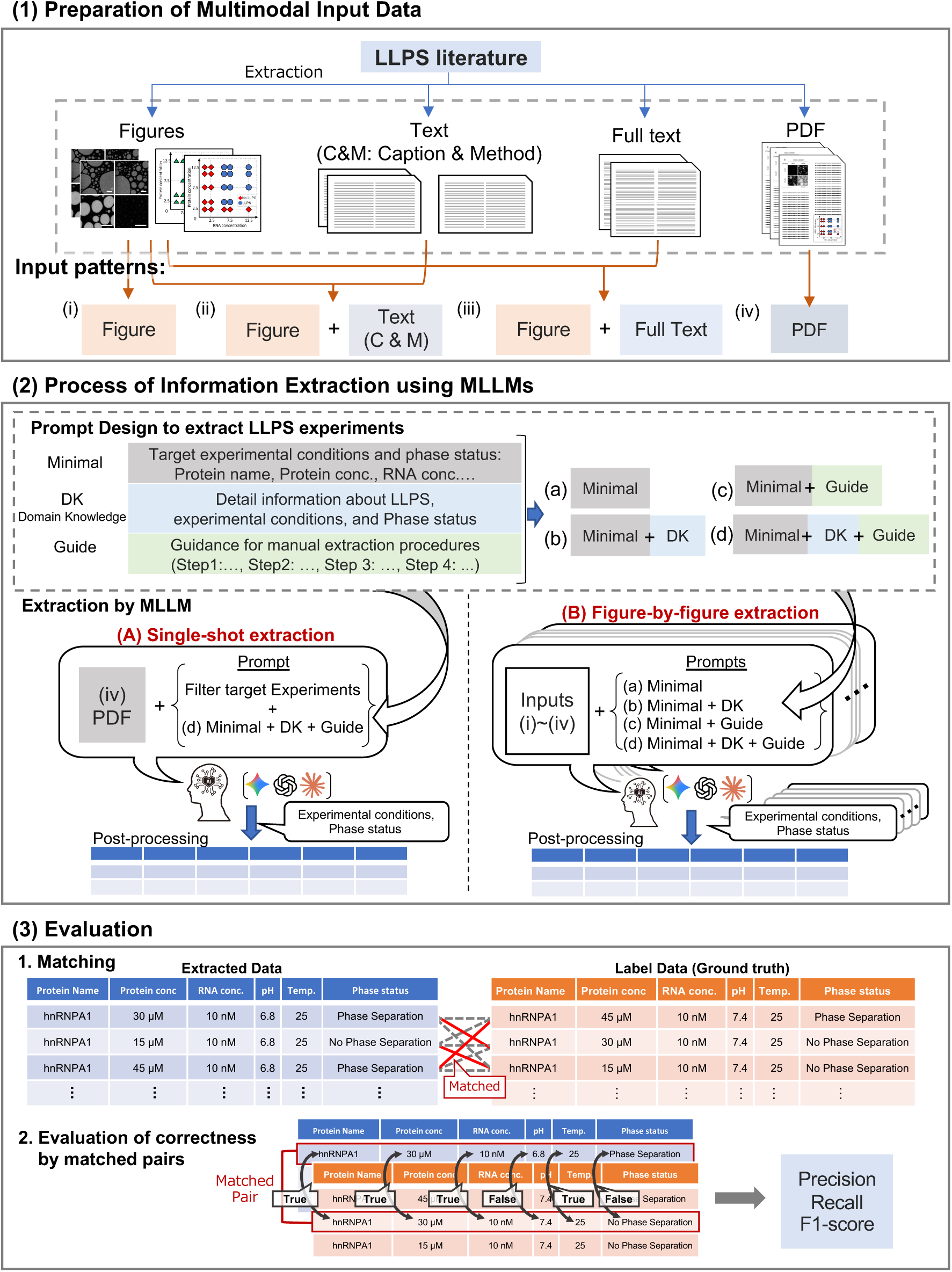
Workflow for Information Extraction and Evaluation of LLPS. In (1), text (full text or captions and methods), images, and PDFs were collected from experimental papers on LLPS, and four input patterns (i)–(iv) were prepared by combining these elements. The examples of microscopic images shown in figures and PDF were reproduced from the paper^40^. In (2), the experimental information (experimental conditions and phase status) is extracted using MLLMs. Four types of prompts (a)–(d) were designed for information extraction, which was performed using two methods: (A) single-shot extraction and (B) figure-by-figure extraction. The extracted data were then organized in tabular form during post-processing. In (3), the extracted results were evaluated. Step (3)-1 processes the matching between extracted and labeled data, and Step (3)-2 shows the evaluation of the matched pairs to calculate correctness and performance metrics.

### 2.2. Evaluation results for all extraction methods

Table 1 shows the F1 scores for each extraction item in the results obtained using the proposed method for 20 papers. Tables S2 and S3 show Precision and Recall. The evaluation results shown here were based on extraction methods using Gemini 2.5 Pro with a temperature set to 0. As a result, figure-by-figure extraction, which sequentially extracts experimental information for each figure, achieved substantially higher F1-scores than the single-shot extraction, which extracts experimental information from a paper in a single process, despite the comparable amounts of extracted data. The method using Figure + Text (C&M: Caption and Method) as the input format and Minimal + DK + Guide as the prompt achieved an average F1-score of 0.836, demonstrating the highest extraction performance among all evaluated methods. Following this, the input patterns Figure (0.809), PDF (0.795), and Figure + Full Text (0.755) using the same Minimal + DK + Guide prompt were ranked. In contrast, the average F1-score for single-shot extraction, without sequential processing, remained at 0.503. As shown in Tables S2 and S3, figure-by-figure extraction consistently outperformed single-shot extraction in both precision and recall, with the Figure + Text (C&M) input and Minimal + DK + Guide prompt achieving the highest scores. Furthermore, when comparing extraction performances across the methods using different prompts, those containing manual extraction guidance showed high performance under experimental conditions that require contextual or coordinate recognition, such as pH, temperature, and concentration. For example, when the input pattern was Figure, adding Guide to the Minimal prompt improved the F1-score of temperature from 0.031 to 0.803 and the F1-score of pH from 0.041 to 0.856. Adding DK to the Minimal prompt significantly affected the target experimental conditions requiring specialized knowledge, such as classification of phase status, with improvements from approximately 0.40 to over 0.70 across all input patterns. Finally, the Minimal + DK + Guide prompt achieved the highest average score across all input patterns.

**Table 1.** Average F1-scores of all proposed methods. The F1-scores were calculated from the results of the single-shot extraction and figure-by-figure extraction using Gemini 2.5 Pro with the temperature parameter set to 0.0. Each column except for “Average” shows the mean F1-scores across all figures (for figure-by-figure extraction) or all papers (for single-shot extraction). The “Average” column shows the overall mean F1-score calculated from the average F1-scores for each field across all figures and papers. The values following “±” indicate the standard deviation (STD) across figures or papers. In the performance metric columns, bold text marks the highest value among all methods. The amount of extracted data is provided as a supplementary information.

| Input | Prompt | Protein name | Protein conc. | RNA conc. | pH | Temperature | Phase Status | Average | Extracted count |
| --- | --- | --- | --- | --- | --- | --- | --- | --- | --- |
| Single-shot Extraction |  |  |  |  |  |  |  |  |  |
| PDF | Minimal + DK + Guide | 0.525 ± 0.284 | 0.464 ± 0.283 | 0.450 ± 0.268 | 0.539 ± 0.306 | 0.539 ± 0.306 | 0.500 ± 0.276 | 0.503 ± 0.270 | 707 |
| Figure-by-figure Extraction |  |  |  |  |  |  |  |  |  |
| Figure | Minimal | 0.693 ± 0.367 | 0.615 ± 0.433 | **0.877 ± 0.163** | 0.041 ± 0.200 | 0.031 ± 0.159 | 0.424 ± 0.384 | 0.447 ± 0.142 | 840 |
|  | Minimal + DK | 0.760 ± 0.325 | 0.615 ± 0.437 | 0.858 ± 0.206 | 0.077 ± 0.261 | 0.052 ± 0.212 | 0.803 ± 0.199 | 0.527 ± 0.180 | 887 |
|  | Minimal + Guide | 0.684 ± 0.374 | 0.808 ± 0.291 | 0.866 ± 0.177 | 0.856 ± 0.214 | 0.803 ± 0.297 | 0.741 ± 0.261 | 0.793 ± 0.180 | 871 |
|  | Minimal + DK + Guide | 0.764 ± 0.315 | 0.800 ± 0.289 | 0.832 ± 0.250 | 0.854 ± 0.214 | 0.794 ± 0.296 | 0.812 ± 0.170 | 0.809 ± 0.187 | 888 |
| Figure + Text<br>(C&M) | Minimal | 0.794 ± 0.267 | 0.824 ± 0.250 | 0.857 ± 0.208 | 0.820 ± 0.268 | 0.346 ± 0.434 | 0.462 ± 0.396 | 0.684 ± 0.182 | 872 |
|  | Minimal + DK | 0.793 ± 0.268 | 0.827 ± 0.257 | 0.843 ± 0.238 | 0.827 ± 0.257 | 0.229 ± 0.383 | 0.811 ± 0.185 | 0.722 ± 0.187 | 893 |
|  | Minimal + Guide | 0.755 ± 0.291 | 0.804 ± 0.268 | 0.836 ± 0.216 | 0.845 ± 0.211 | 0.809 ± 0.269 | 0.563 ± 0.336 | 0.769 ± 0.195 | 857 |
|  | Minimal + DK + Guide | **0.820 ± 0.258** | **0.827 ± 0.254** | 0.863 ± 0.190 | **0.867 ± 0.188** | **0.831 ± 0.255** | 0.810 ± 0.178 | **0.836 ± 0.187** | 848 |
| Figure + Full Text | Minimal | 0.796 ± 0.293 | 0.819 ± 0.277 | 0.789 ± 0.321 | 0.808 ± 0.294 | 0.439 ± 0.451 | 0.390 ± 0.403 | 0.673 ± 0.256 | 925 |
|  | Minimal + DK | 0.785 ± 0.300 | 0.830 ± 0.274 | 0.847 ± 0.246 | 0.811 ± 0.297 | 0.305 ± 0.425 | **0.828 ± 0.208** | 0.734 ± 0.214 | 853 |
|  | Minimal + Guide | 0.750 ± 0.328 | 0.793 ± 0.297 | 0.794 ± 0.304 | 0.813 ± 0.276 | 0.717 ± 0.364 | 0.538 ± 0.362 | 0.734 ± 0.259 | 824 |
|  | Minimal + DK + Guide | 0.724 ± 0.342 | 0.783 ± 0.304 | 0.786 ± 0.305 | 0.803 ± 0.284 | 0.682 ± 0.380 | 0.748 ± 0.277 | 0.755 ± 0.273 | 830 |
| PDF | Minimal | 0.738 ± 0.300 | 0.730 ± 0.341 | 0.809 ± 0.224 | 0.736 ± 0.324 | 0.404 ± 0.442 | 0.426 ± 0.362 | 0.640 ± 0.244 | 712 |
|  | Minimal + DK | 0.754 ± 0.307 | 0.780 ± 0.305 | 0.827 ± 0.231 | 0.791 ± 0.281 | 0.351 ± 0.417 | 0.752 ± 0.217 | 0.709 ± 0.213 | 771 |
|  | Minimal + Guide | 0.766 ± 0.288 | 0.794 ± 0.262 | 0.814 ± 0.217 | 0.830 ± 0.215 | 0.669 ± 0.365 | 0.414 ± 0.359 | 0.715 ± 0.194 | 897 |
|  | Minimal + DK + Guide | 0.757 ± 0.302 | 0.805 ± 0.253 | 0.834 ± 0.191 | 0.837 ± 0.215 | 0.794 ± 0.273 | 0.746 ± 0.206 | 0.795 ± 0.199 | 888 |

To enhance the extraction performance through MLLM family selection and parameter optimization, we conducted additional verification using GPT^56^, Gemini^57^, and Claude^58^ families focusing on the most effective prompt-input data combination: Minimal + DK + Guide prompt with the input pattern Figure + Text (C&M). First, we compared the extraction performances of nine representative MLLMs belonging to three MLLM families, as presented in Method 2.5. As shown in Figure 2, the highest average F1-score for each family was 0.634 for the Claude family (Claude 3.7 Sonnet), 0.807 for the GPT family (GPT-5), and 0.836 for the Gemini family (Gemini 2.5 Pro). Gemini 2.5 Pro achieved the best overall performance, demonstrating its suitability for this extraction task. Next, we optimized the temperature parameter of Gemini 2.5 Pro by testing values of 0.0, 0.1, 0.2, and 0.5 (Table 2). Three independent runs were performed for each temperature, and their averages were used for the evaluation. The results showed the highest average F1-score of 0.836 at a temperature of 0.1, with a maximum F1-score of 0.847. Based on these results, we identified the best-performing method: Gemini 2.5 Pro at a temperature of 0.1, figure-by-figure extraction, input pattern Figure + Text (C&M), and prompt Minimal + DK + Guide. In all subsequent assessments and analyses, we used this best-performing method unless otherwise mentioned. As a supplement, when using the Minimal + DK + Guide prompt with the input pattern Figure + Text (C&M) on Gemini 2.5 Pro, the input token consumption averaged 5,388 tokens. This equates to approximately $0.0072 per figure, or less than $1 for 100 figures (Table S4).

**Figure 2.**
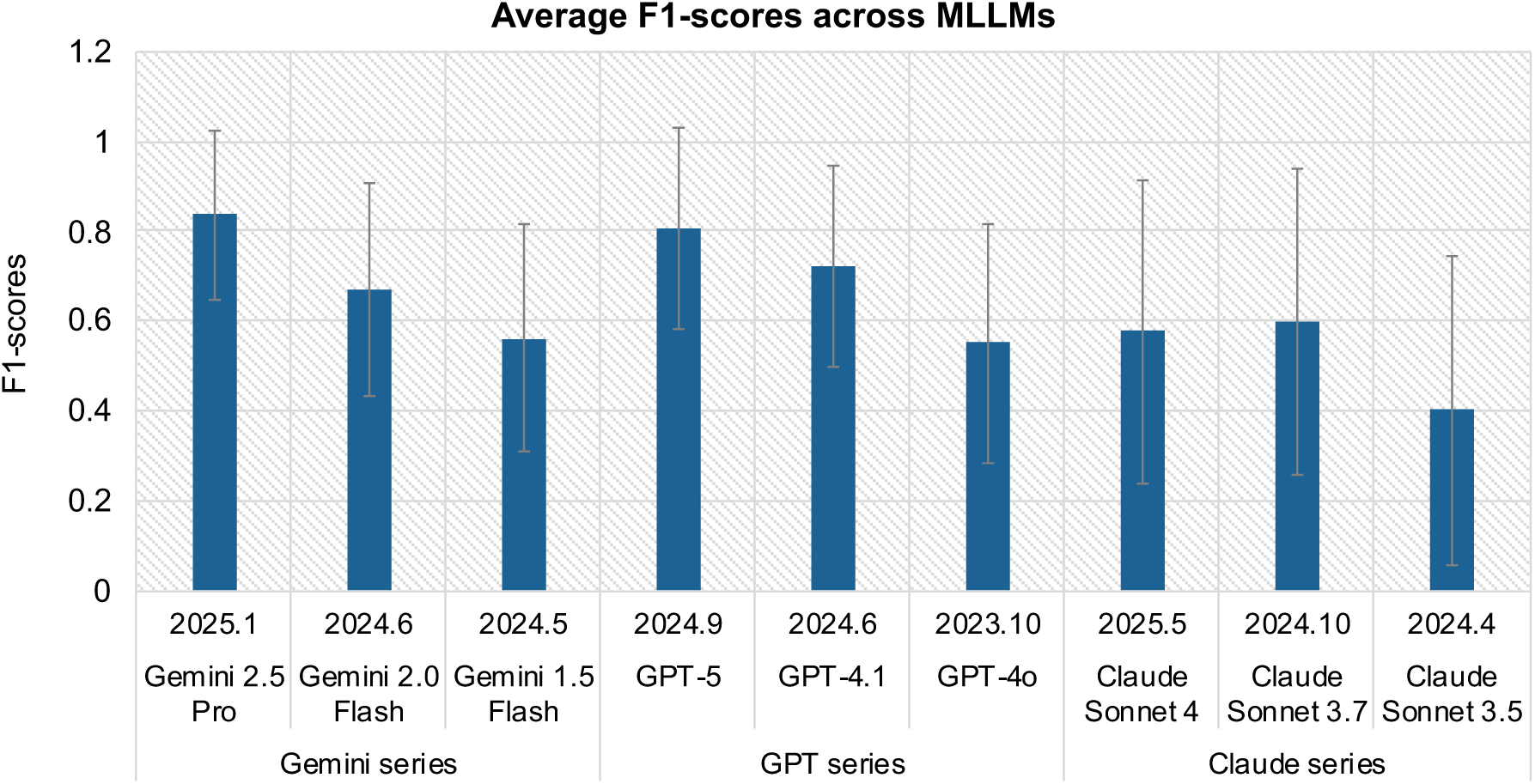
Average F1-scores of different MLLMs for extracting experimental information from LLPS-related figures. Comparison of mean F1-scores across multiple MLLM families under the extraction setting of figure-by-figure extraction, Figure + Text (C&M) as input, and Minimal + Guide + DK as prompt. The date above each MLLM corresponded to its knowledge cutoff date. Error bars indicate the STD of the average F1-score for each figure. Among the tested models, Gemini 2.5 Pro achieved the highest average F1-score, followed by GPT-5 and GPT-4.1. The Claude models showed lower performance, with Claude Sonnet 4 showing the lowest value. These results highlight clear performance differences among the LLM families, with the Gemini and GPT series outperforming the Claude series in this LLPS-specific extraction task.

**Table 2.**
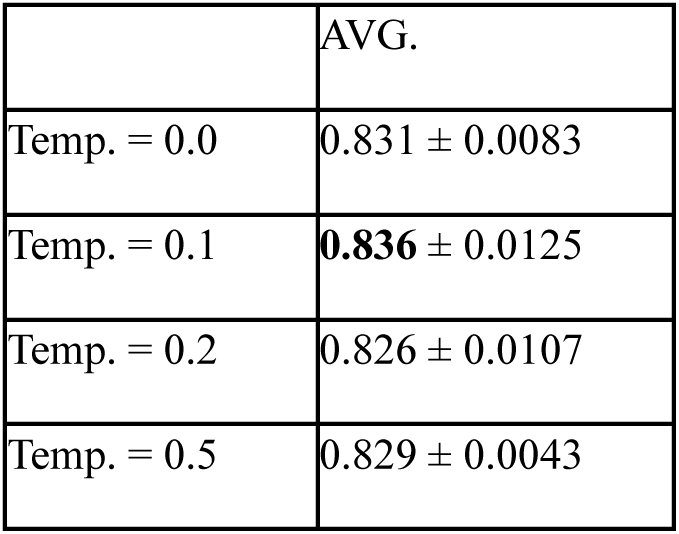
Average F1-scores from temperature parameter optimization for Gemini 2.5 Pro. Comparison of average F1-scores evaluated for Gemini 2.5 Pro across different temperature settings (0.0–0.5) using figure-by-figure extraction, Figure + Text (C&M) input, and Minimal + DK + Guide prompt. Each extraction experiment was repeated three times (N = 3), and the average F1-scores from these repetitions are reported in the “AVG.” column. The values following “±” indicate the STD across the figures.

|  | AVG. |
| --- | --- |
| Temp. = 0.0 | 0.831 ± 0.0083 |
| Temp. = 0.1 | **0.836** ± 0.0125 |
| Temp. = 0.2 | 0.826 ± 0.0107 |
| Temp. = 0.5 | 0.829 ± 0.0043 |

### 2.3. Detailed analysis of extraction performance on scatter plots

To clarify the figure type-specific factors affecting extraction performance, we analyzed the extraction results of the best-performing method for scatter plots and microscope images in the following two sections. First, we focused on scatter plots. As shown in Figure 3A, which presents the extraction performance for each scatter plot, the F1-score exceeded 0.8 for nearly all figures, with an average score of 0.913. Furthermore, for the same input pattern as the best-performing method, the average F1-score was 0.803 for the Minimal prompt, 0.791 for the Minimal + DK prompt, 0.830 for the Minimal + Guide prompt, and 0.848 for the Minimal + DK + Guide prompt, with the prompt containing both DK and Guide showing the highest performance (Table S5).

**Figure 3.**
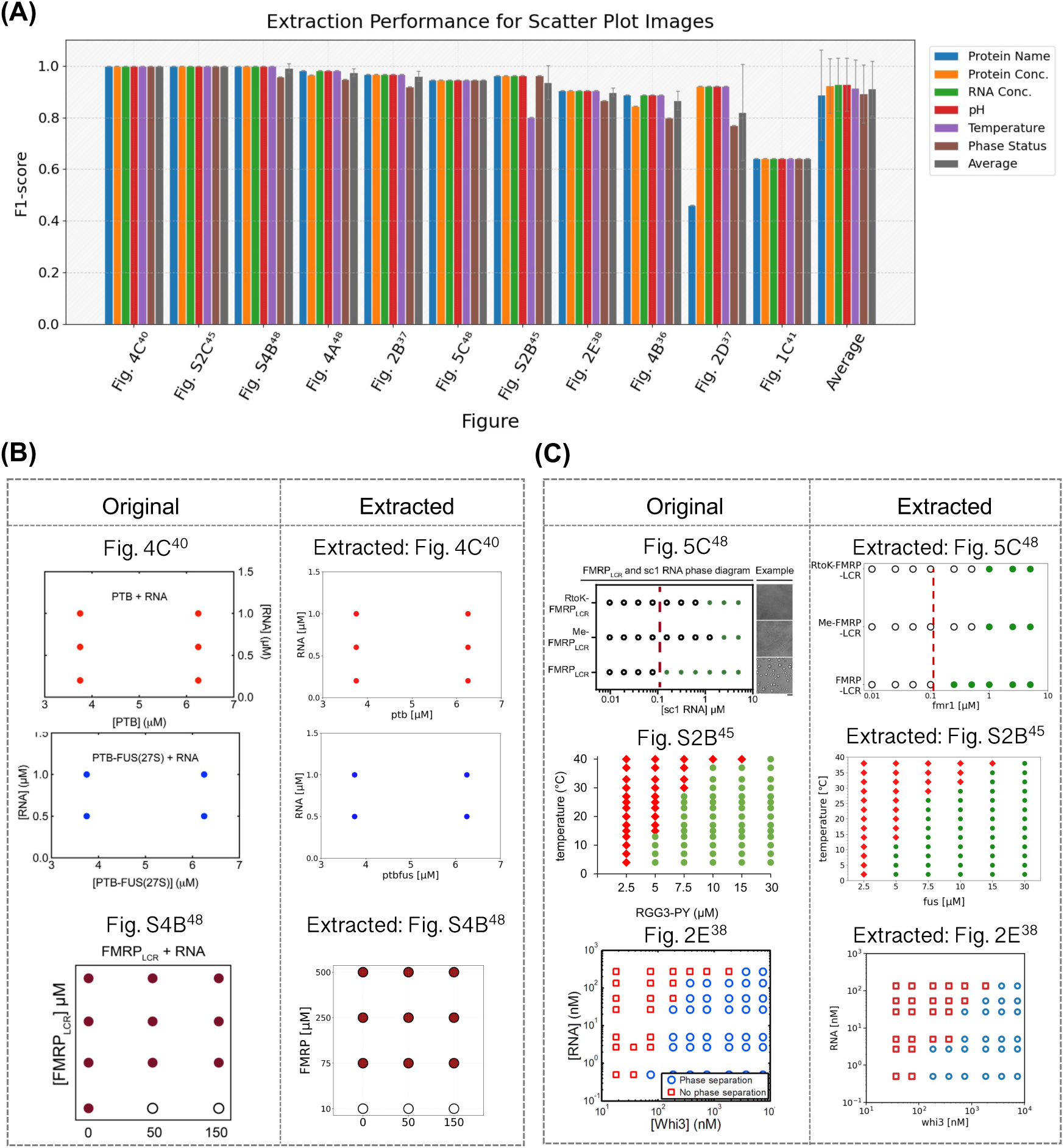
Extraction performance in each scatter plot and comparison between the ground-truth data and extracted data for coordinate extraction. (A) Bar plots show the F1-scores for each figure in the evaluation dataset. Colored bars represent six extracted experimental conditions, comprising protein name, protein concentration, RNA concentration, pH, temperature, and phase status. Gray bars indicate the average F1-score across these items. Error bars represent the STD calculated from the F1-scores for each experimental condition. The rightmost “Average” group shows the overall average F1-scores for each item across all figures, with error bars indicating STD across all figures. (B, C) The left “Original” column shows images reproduced from published articles, and the right column displays the corresponding extraction results. When visualizing data in the “Extracted” figure, the formats of scatter plot elements such as tick markers, labels, marker colors, and sizes were consistent with the corresponding “Original.” Each image is labeled at the top with its reference and subfigure numbers. (B) Examples in which all extracted XY coordinates matched the ground-truth coordinates. (C) Examples in which the extracted XY coordinates contain errors or false positives. The image of Fig. 4C^40^ was reproduced with permission from the paper^40^. The images of Fig. S4B^48^ and Fig. 5C^48^ were reproduced with permission from the paper^48^. The image of Fig. S2B^45^ was reproduced with permission from the paper^45^. The image of Fig. 2E^38^ was reproduced with permission from the paper^38^.

Next, to identify more specific factors, we compared and analyzed successful (Figure 3B) and unsuccessful (Figure 3C) examples of coordinate extraction using actual images. Generally, scatter plots comprise standard elements, such as titles, axis labels, scales, legends, and markers. Most displayed protein and RNA concentrations on the X and Y axes indicated the presence or absence of LLPS in the legend (Figure 3B and 3C). In addition, some featured unique graph structures, including logarithmic scales, categorical variables (Fig. 5C^48^ in Figure 3C), and nonuniform intervals (e.g., Fig. S2B^45^ in Figure 3C). The best-performing method demonstrated high performance in coordinate extraction for scatter plots with few data points, low density, and a regular grid pattern (Figure 3B). However, scatter plots showing errors in coordinate extraction usually contained excessive data points with overlapping regions, leading to false positives or false negatives in those areas (Figure 3C). Furthermore, even with grid-like distributions, errors occurred when the coordinates were slightly offset from the XY grid intersections or when gaps existed in the distribution (Fig. 2E^38^ in Figure 3C). Besides the coordinates, a few errors occurred for the values in the XY-axis labels or titles. For example, Fig. 2E^38^ showed in Figure 3A, an error occurred where only a single protein name was extracted whereas it is a complex protein. Although the performance was slightly reduced for some experimental conditions derived from the text, no major extraction errors directly affecting the average performance of each figure were observed.

### 2.4. Detailed analysis of extraction performance on microscopic images

Here, we analyzed the extraction performance of the best-performing method using 38 microscope images. Figure 4A shows the F1-score for each image and the overall average, with the overall average F1-score achieving a high value of 0.849. However, even with the best-performing method, the extraction performance remained unstable across the figures. Although several figures achieved values close to the F1-score of 1.0, some figures fell below 0.6. For the other prompt conditions, the Minimal prompt yielded 0.660, the Minimal + DK prompt yielded 0.707, and the Minimal + Guide prompt yielded 0.776 (Table S5). Consistent with the results of the scatter plots, the prompt combining DK and Guide performed the best.

**Figure 4.**
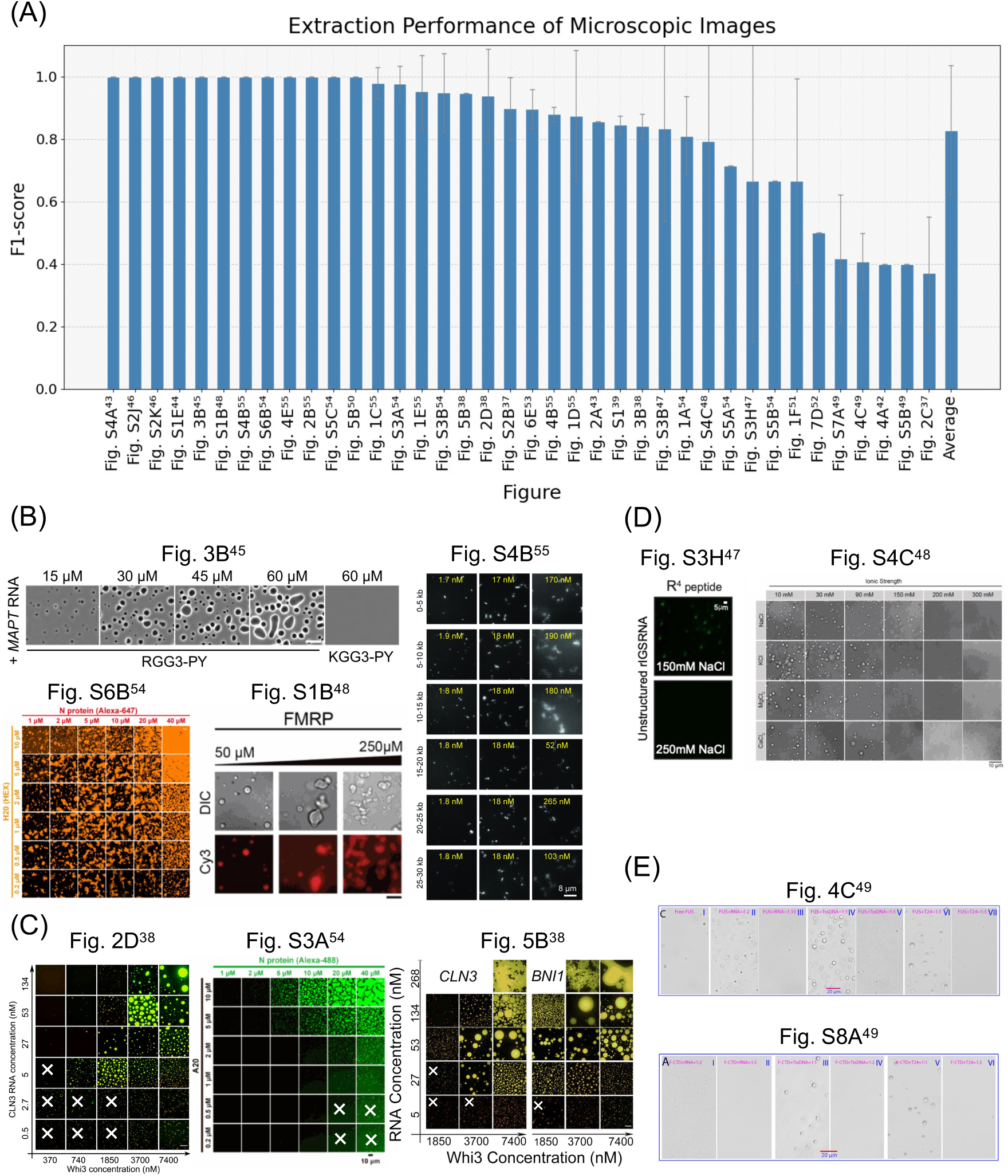
Extraction performance and representative examples for microscope images classified by correctness and error characteristics. (A) Each bar represents the mean F1-score for a single microscopic image, labeled as “Subfigure-number ^Reference-number^.” Error bars, except for the rightmost “Average,” indicate STDs through the experimental conditions. The rightmost bar shows the overall average of the F1-scores, and the error bar represents the STD across all microscopic images. (B) Examples of all extracted experimental conditions were correctly identified. (C) Example of figures showing errors only in the phase status. Experiments marked with a cross (×) indicate a mismatch between the ground truth and extracted value. (D) Examples in which one or more experimental conditions were not extracted (F1-score = 0). (E) Examples with an average F1-score below 0.6. Image of Fig. 4B^45^ was reproduced with permission from the paper^45^. Images of Fig. S6B^54^ and Fig. S3A^54^ were reproduced with permission the paper^54^. Images of Fig. S1B^48^ and Fig. S4C^48^ were reproduced with permission from the paper^48^. Image of Fig.S4B^55^ was reproduced with permission from the paper^55^. Images of Fig. 2D^38^ and Fig. 5B^38^ were reproduced with permission from the paper^38^. Image of Fig. S3H^47^ has been reproduced with permission from the paper^47^. Images of Fig. 4C^49^ and Fig. S8A^49^ were reproduced with permission from the paper^49^.

Next, we analyzed the visual and contextual effects on extraction performance for each experimental condition (Figures 4B–E). We found that the determination of the phase status depends primarily on visual features, specifically the contrast between the granular outlines and the background within the image. Examples with high performance in determining the phase status showed defined droplet formation, sharp outlines, and strong contrast against the background (Figure 4B). Conversely, in examples where errors occurred in the determination of phase status, droplet outlines were unclear, and background contrast was low (Figure 4C). Extraction errors under other experimental conditions were influenced by factors such as the value position, subpanel arrangement, and experimental content. For example, in Figure 4D, critical experimental conditions, such as protein name, concentration, and temperature, were absent from the image itself and scattered throughout the full text, leading to information gaps. Furthermore, examples, such as Figure 4E included experimental conditions outside the scope of this study, such as DNA addition, crowding agents, and time-dependent changes, within the same subfigure, leading to increased false positives and a lower F1-score. In addition, layouts with gaps between panels, particularly figures with numerous subpanels arranged in a grid pattern, was prone to false positives (Fig. 5B^38^ in Figure 4C).

## 3. Discussion

This study showed that the performance of experimental information extraction using MLLMs was significantly influenced by combinations of processing methods, input formats, and prompt contents. Focusing on the differences in performance across the input patterns, Figure + Text (C&M) yielded superior performance compared with the other input patterns. This indicates that the performance can be improved by limiting the input information to data directly relevant to the extraction target, such as figures, captions, and methods sections. Supporting this, the Figure + Full Text series, which combined a single figure with the full text of the target paper, did not yield a higher performance despite the larger information volume. This performance decline could be explained by noise from irrelevant descriptions and formatting errors arising from automatic text extraction from the PDFs. Similarly, the PDF input series, which used both the main and supplementary materials in PDF format, also failed to yield superior performance. This may have been because of structural constraints such as fixed layouts and multi-column formatting, making it difficult to correctly integrate experimental information in different positions. The presence of unrelated sentences and images also induced additional noise. Nevertheless, the PDF input series still achieved F1-scores above 0.80 for several experimental conditions, demonstrating that MLLMs possess high PDF processing capabilities. Finally, whereas the Figure input series achieved moderate performance, the results from the Figure + Text (C&M) input series highlight that adding relevant textual information can further enhance extraction accuracy.

Analysis by figure type showed that the extraction performance was significantly influenced by the prompt content, visual characteristics of the image, and the position of text annotations. Regarding the prompt contents, including Guide or DK within the Minimal prompt improved the extraction performance, with the highest performance achieved when both were combined. Specifically, the Guide was effective for both scatter plots and microscope images, promoting the organization of ambiguous and dispersed information by explicitly showing the extraction procedure. However, DK was effective for microscopic images, encouraging the determination of phase states based on image appearance. For scatter plots, the extraction performance depended on the data point placement and legend clarity, with adequate spacing and explicit labels ensuring high accuracy. In contrast, the overcrowding of data points or gaps in grid-like arrangements led to coordinate extraction errors. For microscope images, the droplet size, contour sharpness, and contrast with the background significantly influenced the phase status extraction. However, current MLLMs often fail to perform adequately and produce erroneous outputs when dealing with low-visibility experimental images that require contextual judgment. Thus, whereas scatter plots enable relatively stable extraction supported by visual cues, microscopic images require integrating both visual information and the context of the phases, leading to unstable performance. Furthermore, for both image types, the methods section often lists numerous test conditions that could cause erroneous extraction of unrelated information.

The processing method and prompts proposed in this study demonstrated a certain level of performance; however, challenges remain in both the extraction workflow and performance for practical implementation. From a workflow perspective, the proposed method requires manual intervention for figure selection and extraction of captions and methodologies. Although several automated PDF extraction tools have been proposed, they often result in text corruption, image quality degradation, and loss of the spatial relationship between figures and text^59,60^. Currently, manual processing is the most reliable method. Furthermore, processing all images without relevance filtering increases the computational cost and risks, generating numerous false positives and negatives.

However, distinguishing target figures from nontarget figures using MLLMs requires integrated interpretation of image information and text descriptions of methods and procedures, yet current models show insufficient classification performance for this task (F1-score = 0.44, Figure S1A). Numerous misclassifications occurred in cases where images appeared to be the target when viewed alone, whereas careful review of the content revealed that they were non-targets (Figure S1B). Therefore, although the proposed method shows promising performance, significant challenges remain for full automation.

From the extraction performance perspective, the results are not of sufficient quality for direct use in database construction or downstream analyses. One reason is that, compared to materials science and medicine, where information extraction has been extensively studied, the LLPS domain faces greater technical challenges because of the frequent occurrence of unstructured, multi-parameter, and ambiguous expressions. In materials science, entity- and relation-extraction tasks are relatively well defined (typically targeting compounds, properties, and synthesis steps), with multiple studies achieving high performance using LLM^14–16^. In contrast, LLPS requires the integration of multiple heterogeneous elements, including numerical values, such as temperature and concentration, protein names, and the presence or absence of phase separation, into a coherent experimental context. In particular, for microscopic images, clear ground-truth labels rarely exist, requiring the interpretation of both images and context. Similarly, scatter plots must treat each data point as a single experiment and reconstruct the values of the experimental conditions from the axis labels, titles, and legends. These tasks differ from traditional named entity recognition and relation extraction, requiring multimodal processing that is distinct from the existing domains. Therefore, achieving a performance equivalent to that of these fields remains difficult. However, as MLLMs are rapidly advancing and have shown notable improvements in this task (Figure 2), the issues identified here may naturally be mitigated through their continued evolution. Given that MLLM usage costs are trending downward, data collection will become more accurate and cost-efficient.

## 4. Conclusion

In this study, we proposed extraction methods for LLPS experimental information using MLLMs and evaluated the effects of input formats and prompt design on extraction performance. The combination of figures with captions and method sections together with structured prompts incorporating DK achieved the highest performance in extracting experimental information. Furthermore, an analysis of the influence of visual features on performance revealed that scatter plots with simple and clear structures enabled highly accurate extraction, with over 70% of the figures achieving an F1-score above 0.9. In contrast, dense point distributions and unclear axes or legends are the primary sources of error, reducing the performance of such images. Microscopic images also showed relatively high performance; however, low droplet visibility and the presence of nontarget conditions were identified as factors that reduced the extraction accuracy. Nevertheless, the extraction performance of the current MLLMs is not yet sufficiently reliable for direct use in database construction or large-scale analyses, and challenges remain for full automation. Experiments with multiple conditions, ambiguous microscopy images, and densely populated scatter plots remain particularly difficult to extract accurately. However, the performance of MLLMs has been improving rapidly, with recent models exhibiting remarkable advances in logical reasoning and image understanding. Therefore, the challenges identified in this study are expected to be resolved through ongoing technological progress, including model advancement, dataset expansion, and processing automation. Although this study focused on LLPS, descriptions integrating visual and textual information are common across numerous fields of biology. Therefore, the findings of this study provide insight into the extraction of diverse biological phenomena.

## 5. Methods

### 5.1. Preparation of multimodal input data

To verify the effect of the input data format on the extraction performance, we prepared the following input patterns, as shown in (1) of Figure 1: (i) Figure only, (ii) Figure + Text (C&M: Captions and Methods), (iii) Figure + Full Text, and (iv) PDF. We focused on microscopy images and scatter plots because LLPS studies primarily visualize experimental observations in this way. The highest-resolution images were downloaded individually for each subfigure. Text (C&M) was collected manually from relevant sections of the target papers. The full text was automatically extracted from the main text and supplementary PDFs using PyMuPDF^61^. PDFs were downloaded from the publisher’s website. With input patterns (i) to (iv), we evaluated how different input formats affect the extraction results and examined whether separating figures and text from PDF documents is necessary.

### 5.2. Process of information extraction using MLLMs

In this study, we aimed to extract information about LLPS experiments noted in scatter plots and microscopy images, focusing on six commonly recorded experimental conditions: protein name, protein concentration, RNA concentration, temperature, buffer pH, and phase status. To achieve more advanced extraction performance, we compared combinations of multiple input formats, various prompt designs, and two processing methods. For the input formats, we used (i)–(iv) created in Section 2.1. For prompts, we designed the following four types with differing structures and contents, as shown in the upper half of (2) in Figure 1: (a) Minimal, a prompt containing the target experimental conditions and brief extraction instructions; (b) Minimal + DK, a prompt comprising the Minimal prompt with a detailed explanation (DK) for each item; (c) Minimal + Guide, a prompt comprising the Minimal prompt with a manual extraction procedure guide (Guide) provided by an expert; and (d) Minimal + DK + Guide, a prompt comprising the Minimal prompt with both DK and Guide added. Details of each prompt are provided in the sections 1-4 of the supplementary material “Additional_file2_prompts.docx.” Two main processing methods were adopted and experimented with. The first method, single-shot extraction, extracted all the experimental information from the entire paper without referencing specific figures (Figure 1, left panel: single-shot extraction). This method uses a prompt combining filter statements containing the extraction criteria for target experiments with the prompt (d) Minimum + DK + Guide and extracts experimental information from (iv) PDF. The prompt is provided in the section 5 of the supplementary material “Additional_file2_prompts.docx.” In the second method, figure-by-figure extraction, each figure was treated as a single processing unit, and the experimental information corresponding to the specified figures was sequentially extracted (Figure 1, right panel: figure-by-figure extraction). This method verified 16 different configurations for extracting experimental information from input formats (i)– (iv) using prompts (a)–(d). The figure specification within the processing units is typically based on subfigure numbers. When multiple scatter plots existed within a single subfigure, their respective positions (right, left, top, bottom, etc.) were specified explicitly.

After the extraction processes, the textual output extracted by the MLLMs was formatted into a tabular structure, and the representation of each extracted value was standardized using the following method. For the numerical experimental conditions, including temperature, RNA concentration, and protein concentration, the units were normalized using predefined rule-based procedures. Temperature values were converted to Celsius, with “RT” and room temperature uniformly converted to “25 °C.” RNA and protein concentrations were standardized to “μM.” Values with units such as “mg/mL,” which cannot be converted to molar concentrations without information on molecular weight or solvent volume, were used as is. Protein names were standardized to ensure consistent notation for the same protein.

### 5.3. Evaluation for information extraction

As shown in (3)-1 of Figure 1, the order of the extracted data and label data do not necessarily match. Therefore, establishing a correspondence between the extracted and label data is crucial. To achieve this, we formulated the matching of the extracted data and label data as an assignment problem and solved it using the Hungarian method^62^. This is a method for efficiently determining the minimum cost matching based on a cost matrix. In this study, the rows and columns of the cost matrix correspond to the extracted and label data, respectively. The elements of the cost matrix represent the costs computed from pairs of extracted data and label data, with each cost indicating the average distance across all experimental conditions. For the numerical fields of the experimental conditions, including RNA concentration, protein concentration, pH, and temperature, each distance was defined as |*x_pred_* − *x_true_*|/(*x_max_* − *x_min_*), where *x_pred_* is the value of the extracted data, *x_pred_* is the value of the label data, and *x_max_* and *x_min_* are the maximum and minimum values of the label data in the evaluated figure. For the textual field of the experimental condition, the distance was defined as 1 − *s*, where *s* is the similarity score from Sequence Matcher^63^, with *s* = 1 for an exact match. Matching was performed on a per-paper basis for single-shot extraction, and on a per-figure basis for figure-by-figure extraction.

Next, as shown in (3)-2 of Figure 1, we calculated evaluation metrics based on the distance between each experimental condition for the matched data pairs identified by the Hungarian algorithm. For the five experimental conditions, except for the binary field of the phase status, each distance was calculated using the same method as that used in the matching process. In the phase status, which is a binary field indicating the presence or absence of LLPS, the distance was set to 0 for a match and 1 for a mismatch. The matched extracted data were counted as true positives if their distance was at most 0.2; otherwise, they were counted as false positives. Data without a corresponding pair were treated as follows: extracted data without matched label data were counted as false positives, and label data without matched extracted data were counted as false negatives. Based on the number of true positives, false positives, and false negatives, we calculated the precision, recall, and F1-scores. In figure-by-figure extraction, this metrics were computed for each figure. In the single-shot extraction, they were computed for each paper. Finally, the average scores across all papers in the single-shot extraction and across all figures in the figure-by-figure extraction were used.

### 5.4. Evaluation dataset for extraction

The evaluation of the extraction methods used a curated dataset constructed from 20 experimental papers on LLPS^36–55^ registered with RNAPSEC^30^. This curated dataset comprises reliable labeled data, as the original images and corresponding manually annotated experimental information are available in RNAPSEC. The references for each study and their corresponding reference numbers are listed in Table S1. The following requirements were selected for extraction: experiments directly observing LLPS; use of a single type of protein and RNA (fusion proteins were counted as a single type); experiments varying only protein, RNA, pH, salt, and temperature, as shown through microscope images or scatter plots; and all experimental conditions subject to extraction explicitly stated or implied within the paper. In total, 904 experiments derived from 49 LLPS experimental figures from 20 papers^36–55^ were selected.

### 5.5. MLLMs setting for extraction

MLLMs used in this study included Gemini 2.5 Pro (gemini-2.5-pro-preview-06-05), Gemini 2.0 Flash (gemini-2.0-flash), and Gemini 1.5 Flash (gemini-1.5-flash) provided by Google; Claude sonnet 4 (claude-sonnet-4-20250514), Claude sonnet 3.5 (claude-3.5-sonnet-20241022), and Claude sonnet 3.7 (claude-3.7-sonnet-20250219) provided by Anthropic; GPT-5 (gpt-5), GPT-4.1 (gpt-4.1-2025-04-14), and GPT-4o (gpt-4o-2024-08-06) provided by OpenAI. For input pattern (iv) PDF, each model was accessed through its official API provided by the respective developer. For the other input patterns, MLLM access was implemented via LangChain^64^, which provides standardized interfaces for the official APIs of each model. In experiments where no specific model parameters were mentioned, the temperature parameter was set to 0.0, and the next token with the highest probability was selected. All other parameters were set to default values. In experiments optimizing the temperature parameter, values of 0.0, 0.1, 0.2, and 0.5 were examined.

## Supporting information

Supporting Information (Prompts)

## Abbreviations

LLPS: Liquid-liquid phase separation
MLLM: Multimodal large language model
DK: Domain knowledge
C & M: Caption and methods
STD: Standard deviation

## Data availability

Extraction models are available in the GitHub repository (https://github.com/ycu-iil/MLLMIE).

## Acknowledgements

This work was supported by the JST FOREST program (Grant Number: JPMJFR232U) and JST BOOST program (Grant Number: JPMJBY24F0). Additional support was provided by the Ministry of Education, Culture, Sports, Science, and Technology (MEXT) under grant Data Creation and Utilization Type Material Research and Development Project (Grant Number: JPMXP1122683430), Simulation and AI-driven Next-Generation Medicine and Drug Discovery based on “Fugaku” (Grant Number: JPMXP1020230120), and feasibility studies for the next-generation computing infrastructure.

## Author’s contributions

KC, SI, and KT designed the research concept. KC performed the programming, data collection, and manuscript writing. SF, SI, and KT edited and verified the manuscript and data, and supervised the project. All the authors have reviewed and approved the final manuscript.

## Supplementary information

The supplementary tables and figure are available at “Additional_file1_SI.docx.” The prompts used in this study are provided in “Additional_file2_prompts.docx.”

