## Supporting Information (Tables and Figure) for "Assessing the Performance of LLMs in Multimodal Information Extraction for Biological Research: A Case Study on LLPS"

Ka Yin Chin^1^, Satoru Fujii^1^, Shoichi Ishida^1^, and Kei Terayama^1,2,3,4*^

^1^ Graduate School of Medical Life Science, Yokohama City University, 1-7-29, Suehiro-cho, Tsurumi-ku, Yokohama, Kanagawa, 230-0045, Japan.

^2^ RIKEN Center for Advanced Intelligence Project, 1-4-1, Nihonbashi, Chuo-ku, Tokyo 103-0027, Japan.

^3^ MDX Research Center for Element Strategy, Tokyo Institute of Technology, 4259 Nagatsuta-cho, Midori-ku, Yokohama, Kanagawa, 226-8501, Japan.

^4^Department of Life Science and Technology, School of Life Science and Technology, Institute of Science Tokyo, 4259 Nagatsuta-cho, Midori-ku, Yokohama, Kanagawa, 226-8501, Japan.

^*^

**Contents:**

Table S1. Correspondence between papers and reference numbers in the evaluation dataset.

Table S2. Precision scores of all proposed methods.

Table S3. Recall scores of all proposed methods.

Table S4. Average and total extraction costs per figure for each MLLMs.

Table S5. Comparison of extraction performance between scatter plots and microscopic images.

Figure S1. F1-scores and failure examples in extracting target figure numbers.

**Table S1. Correspondence between papers and reference numbers in the evaluation dataset.** The table lists the papers used in the evaluation and their corresponding reference numbers in the main text.

| Reference number | Reference |
| --- | --- |
| 36 | Molliex, A. et al. Phase separation by low complexity domains promotes stress granule assembly and drives pathological fibrillization. *Cell* **163**, 123–133 (2015). 10.1016/j.cell.2015.09.015, PubMed: 26406374. |
| 37 | Lin, Y., Protter, D. S. W., Rosen, M. K. & Parker, R. Formation and maturation of phase-separated liquid droplets by RNA-binding proteins*. Mol. Cell* **60**, 208–219 (2015). 10.1016/j.molcel.2015.08.018, PubMed: 26412307. |
| 38 | Zhang, H. et al*.* RNA controls PolyQ protein phase transitions. *Mol. Cell* **60**, 220–230 (2015). 10.1016/j.molcel.2015.09.017, PubMed: 26474065. |
| 39 | Smith, J. et al. Spatial patterning of P granules by RNA-induced phase separation of the intrinsically disordered protein MEG-3 . *eLife* **5**, e21337 (2016). 10.7554/eLife.21337, PubMed: 27914198. |
| 40 | Lin, Y., Currie, S. L. & Rosen, M. K. Intrinsically disordered sequences enable modulation of protein phase separation through distributed tyrosine motifs. *J. Biol. Chem.* **292**, 19110–19120 (2017). 10.1074/jbc.M117.800466, PubMed: 28924037. |
| 41 | Wei, M. T. et al. Phase behaviour of disordered proteins underlying low density and high permeability of liquid organelles. *Nat. Chem.* **9**, 1118–1125 (2017). 10.1038/nchem.2803, PubMed: 29064502. |
| 42 | Protter, D. S. W. et al. Intrinsically disordered regions can contribute promiscuous interactions to RNP granule assembly. *Cell Rep.* **22**, 1401–1412 (2018). 10.1016/j.celrep.2018.01.036, PubMed: 29425497. |
| 43 | Maharana, S. et al. RNA buffers the phase separation behavior of prion-like RNA-binding proteins . *Science* **360**, 918–921 (2018). 10.1126/science.aar7366, PubMed: 29650702. |
| 44 | Langdon, E. M. et al. mRNA structure determines specificity of a polyQ-driven phase separation. *Science* **360**, 922–927 (2018). 10.1126/science.aar7432, PubMed: 29650703. |
| 45 | Hofweber, M. et al. Phase separation of FUS is suppressed by its nuclear import receptor and arginine methylation. *Cell* **173**, 706–719.e13 (2018). 10.1016/j.cell.2018.03.004, PubMed: 29677514. |
| 46 | Kroschwald, S. et al. Different material states of Pub1 condensates define distinct modes of stress adaptation and recovery. *Cell Rep.* **23**, 3327–3339 (2018). 10.1016/j.celrep.2018.05.041, PubMed: 29898402. |
| 47 | Wang, M. et al*.* Stress-induced low complexity RNA activates physiological amyloidogenesis. *Cell Rep.* **24**, 1713–1721.e4 (2018). 10.1016/j.celrep.2018.07.040, PubMed: 30110628. |
| 48 | Tsang, B. et al. Phosphoregulated FMRP phase separation models activity-dependent translation through bidirectional control of mRNA granule formation. *Proc. Natl Acad. Sci. U. S. A.* **116**, 4218–4227 (2019). 10.1073/pnas.1814385116, PubMed: 30765518. |
| 49 | Kang, J., Lim, L., Lu, Y. & Song, J. A unified mechanism for LLPS of ALS/FTLD-causing FUS as well as its modulation by ATP and oligonucleic acids. *PLOS Biol.* **17**, e3000327 (2019). 10.1371/journal.pbio.3000327, PubMed: 31188823. |
| 50 | Tari, M. et al. U2AF^65^ assemblies drive sequence-specific splice site recognition. *EMBO Rep.* **20**, e47604 (2019). 10.15252/embr.201847604, PubMed: 31271494. |
| 51 | Ries, R. J. et al. m6A enhances the phase separation potential of mRNA. *Nature* **571**, 424–428 (2019). 10.1038/s41586-019-1374-1, PubMed: 31292544. |
| 52 | Niaki, A. G. et al*.* Loss of dynamic RNA interaction and aberrant phase separation induced by two distinct types of ALS/FTD-linked FUS mutations. *Mol. Cell* **77**, 82–94.e4 (2020). 10.1016/j.molcel.2019.09.022, PubMed: 31630970. |
| 53 | Huo, X. et al*.* The nuclear matrix protein SAFB cooperates with major satellite RNAs to stabilize heterochromatin architecture partially through phase separation. *Mol. Cell* **77**, 368–383.e7 (2020). 10.1016/j.molcel.2019.10.001, PubMed: 31677973. |
| 54 | Chen, H. et al*.* Liquid–liquid phase separation by SARS-CoV-2 nucleocapsid protein and RNA. *Cell Res.* **30**, 1143–1145 (2020). 10.1038/s41422-020-00408-2, PubMed: 32901111. |
| 55 | Jack, A. et al*.* SARS-CoV-2 nucleocapsid protein forms condensates with viral genomic RNA. *PLOS Biol.* **19**, e3001425 (2021). 10.1371/journal.pbio.3001425, PubMed: 34634033. |

**Table S2. Precision scores of all proposed methods.** The precision scores were calculated from the results of the single-shot extraction and figure-by-figure extraction using Gemini 2.5 Pro with the temperature parameter set to 0.0, as shown in Table 1. Each column, except for “Average,” shows the mean precision across all figures (for figure-by-figure extraction) or all papers (for single-shot extraction). The “Average” column shows the overall mean precision calculated from the average precision values for each field across all figures and papers. The values following “±” indicate the standard deviation (STD) across figures or papers. In the performance metric columns, bold text marks the highest value among all methods. The amount of extracted data is provided as a supplementary information. Abbreviations: conc., concentration.

| Input | Prompt | Protein name | Protein conc. | RNA conc. | pH | Temperature | Phase Status | Average | Extracted count |
| --- | --- | --- | --- | --- | --- | --- | --- | --- | --- |
| Single-shot Extraction | | | | | | | | | |
| PDF | Minimal + DK + Guide | 0.628± 0.340 | 0.492 ± 0.318 | 0.516 ± 0.367 | 0.595 ± 0.381 | 0.595 ± 0.380 | 0.606 ± 0.339 | 0.572 ± 0.307 | 707 |
| Figure-by-figure Extraction | | | | | | | | | |
| Figure | Minimal | 0.705 ± 0.392 | 0.628 ± 0.452 | 0.882 ± 0.209 | 0.041 ± 0.200 | 0.034 ± 0.169 | 0.432 ± 0.408 | 0.453 ± 0.170 | 840 |
|  | Minimal + DK | 0.755 ± 0.349 | 0.618 ± 0.450 | 0.850 ± 0.256 | 0.075 ± 0.257 | 0.056 ± 0.223 | 0.795 ± 0.252 | 0.525 ± 0.204 | 887 |
|  | Minimal + Guide | 0.683 ± 0.392 | 0.800 ± 0.317 | 0.859 ± 0.228 | 0.851 ± 0.258 | 0.796 ± 0.327 | 0.729 ± 0.296 | 0.786 ± 0.220 | 871 |
|  | Minimal + DK + Guide | 0.758 ± 0.337 | 0.791 ± 0.319 | 0.827 ± 0.285 | 0.838 ± 0.262 | 0.781 ± 0.327 | 0.802 ± 0.227 | 0.799 ± 0.229 | 888 |
| Figure + Text (C&M) | Minimal | 0.796 ± 0.300 | 0.823 ± 0.286 | 0.857 ± 0.251 | 0.816 ± 0.302 | 0.354 ± 0.453 | 0.457 ± 0.406 | 0.684 ± 0.214 | 872 |
|  | Minimal + DK | 0.793 ± 0.296 | 0.824 ± 0.292 | 0.846 ± 0.268 | 0.826 ± 0.292 | 0.219 ± 0.380 | 0.810 ± 0.231 | 0.720 ± 0.220 | 893 |
|  | Minimal + Guide | 0.750 ± 0.326 | 0.799 ± 0.309 | 0.832 ± 0.266 | 0.841 ± 0.262 | 0.807 ± 0.310 | 0.546 ± 0.356 | 0.763 ± 0.239 | 857 |
|  | Minimal + DK + Guide | **0.832 ± 0.289** | **0.835 ± 0.289** | 0.871 ± 0.233 | **0.876 ± 0.233** | **0.842 ± 0.289** | **0.819 ± 0.226** | **0.846 ± 0.232** | 848 |
| Figure + Full Text | Minimal | 0.777 ± 0.319 | 0.798 ± 0.306 | 0.779 ± 0.342 | 0.790 ± 0.324 | 0.427 ± 0.449 | 0.382 ± 0.402 | 0.659 ± 0.276 | 925 |
|  | Minimal + DK | 0.784 ± 0.321 | 0.830 ± 0.303 | 0.845 ± 0.279 | 0.811 ± 0.324 | 0.305 ± 0.433 | 0.826 ± 0.246 | 0.734 ± 0.245 | 853 |
|  | Minimal + Guide | 0.765 ± 0.353 | 0.807 ± 0.325 | 0.814 ± 0.326 | 0.827 ± 0.304 | 0.734 ± 0.388 | 0.538 ± 0.380 | 0.748 ± 0.286 | 824 |
|  | Minimal + DK + Guide | 0.731 ± 0.369 | 0.787 ± 0.338 | 0.795 ± 0.335 | 0.808 ± 0.319 | 0.684 ± 0.404 | 0.755 ± 0.314 | 0.760 ± 0.308 | 830 |
| PDF | Minimal | 0.828 ± 0.296 | 0.787 ± 0.345 | **0.911 ± 0.194** | 0.817 ± 0.337 | 0.449 ± 0.479 | 0.470 ± 0.409 | 0.710 ± 0.228 | 712 |
|  | Minimal + DK | 0.816 ± 0.322 | 0.820 ± 0.321 | 0.882 ± 0.243 | 0.847 ± 0.299 | 0.386 ± 0.459 | 0.807 ± 0.244 | 0.760 ± 0.229 | 771 |
|  | Minimal + Guide | 0.811 ± 0.313 | 0.833 ± 0.283 | 0.854 ± 0.244 | 0.870 ± 0.239 | 0.705 ± 0.393 | 0.437 ± 0.396 | 0.752 ± 0.226 | 897 |
|  | Minimal + DK + Guide | 0.788 ± 0.321 | 0.828 ± 0.279 | 0.859 ± 0.211 | 0.859 ± 0.240 | 0.814 ± 0.292 | 0.769 ± 0.243 | 0.820 ± 0.225 | 888 |

**Table S3. Recall scores of all proposed methods.** The recall scores were calculated from the results of the single-shot extraction and figure-by-figure extraction using Gemini 2.5 Pro with the temperature parameter set to 0.0, as shown in Table 1. Each column, except for “Average,” shows the mean recall across all figures (for figure-by-figure extraction) or all papers (for single-shot extraction). The “Average” column shows the overall mean recall calculated from the average recall values for each field across all figures and papers. The values following “±” indicate the STD across figures and papers. In the performance metric columns, bold text marks the highest value among all methods. The amount of extracted data is provided as supplemental information. Abbreviations: conc., concentration.

| Input | Prompt | Protein name | Protein conc. | RNA conc. | pH | Temperature | Phase Status | Average | Extracted count |
| --- | --- | --- | --- | --- | --- | --- | --- | --- | --- |
| Single-shot Extraction | | | | | | | | | |
| PDF | Minimal + DK + Guide | 0.714 ± 0.369 | 0.627 ± 0.377 | 0.610 ± 0.333 | 0.733 ± 0.381 | 0.732 ± 0.381 | 0.685 ± 0.350 | 0.684 ± 0.339 | 707 |
| Figure-by-figure Extraction | | | | | | | | | |
| Figure | Minimal | 0.725 ± 0.375 | 0.639 ± 0.448 | 0.927 ± 0.148 | 0.041 ± 0.200 | 0.029 ± 0.154 | 0.449 ± 0.401 | 0.468 ± 0.133 | 840 |
|  | Minimal + DK | 0.821 ± 0.329 | 0.652 ± 0.455 | 0.945 ± 0.138 | 0.080 ± 0.271 | 0.049 ± 0.207 | 0.889 ± 0.157 | 0.573 ± 0.154 | 887 |
|  | Minimal + Guide | 0.729 ± 0.389 | 0.870 ± 0.292 | 0.937 ± 0.142 | 0.924 ± 0.190 | 0.870 ± 0.291 | 0.813 ± 0.265 | 0.857 ± 0.157 | 871 |
|  | Minimal + DK + Guide | 0.818 ± 0.329 | 0.865 ± 0.291 | 0.892 ± 0.236 | 0.933 ± 0.180 | 0.869 ± 0.291 | 0.887 ± 0.152 | 0.877 ± 0.156 | 888 |
| Figure + Text (C&M) | Minimal | 0.845 ± 0.262 | 0.880 ± 0.245 | 0.913 ± 0.192 | 0.878 ± 0.263 | 0.367 ± 0.461 | 0.495 ± 0.415 | 0.730 ± 0.185 | 872 |
|  | Minimal + DK | 0.852 ± 0.271 | 0.901 ± 0.239 | 0.900 ± 0.214 | 0.901 ± 0.238 | 0.269 ± 0.435 | 0.884 ± 0.162 | 0.784 ± 0.170 | 893 |
|  | Minimal + Guide | 0.821 ± 0.292 | 0.874 ± 0.261 | 0.906 ± 0.194 | 0.916 ± 0.187 | 0.875 ± 0.260 | 0.636 ± 0.370 | 0.838 ± 0.193 | 857 |
|  | Minimal + DK + Guide | 0.868 ± 0.257 | 0.889 ± 0.243 | 0.925 ± 0.163 | 0.929 ± 0.157 | 0.890 ± 0.243 | 0.871 ± 0.165 | 0.895 ± 0.161 | 848 |
| Figure + Full Text | Minimal | 0.865 ± 0.283 | 0.894 ± 0.263 | 0.839 ± 0.314 | 0.882 ± 0.283 | 0.478 ± 0.485 | 0.412 ± 0.416 | 0.728 ± 0.241 | 925 |
|  | Minimal + DK | 0.831 ± 0.309 | 0.884 ± 0.262 | 0.906 ± 0.228 | 0.866 ± 0.290 | 0.329 ± 0.453 | 0.886 ± 0.188 | 0.784 ± 0.202 | 853 |
|  | Minimal + Guide | 0.796 ± 0.335 | 0.844 ± 0.303 | 0.828 ± 0.304 | 0.865 ± 0.278 | 0.763 ± 0.378 | 0.596 ± 0.400 | 0.782 ± 0.266 | 824 |
|  | Minimal + DK + Guide | 0.780 ± 0.355 | 0.853 ± 0.299 | 0.846 ± 0.300 | 0.874 ± 0.272 | 0.753 ± 0.394 | 0.813 ± 0.270 | 0.820 ± 0.264 | 830 |
| PDF | Minimal | 0.746 ± 0.329 | 0.752 ± 0.364 | 0.815 ± 0.269 | 0.747 ± 0.351 | 0.403 ± 0.455 | 0.434 ± 0.382 | 0.650 ± 0.273 | 712 |
|  | Minimal + DK | 0.758 ± 0.333 | 0.813 ± 0.321 | 0.853 ± 0.254 | 0.816 ± 0.302 | 0.364 ± 0.438 | 0.778 ± 0.245 | 0.730 ± 0.233 | 771 |
|  | Minimal + Guide | 0.796 ± 0.309 | 0.848 ± 0.265 | 0.872 ± 0.219 | 0.887 ± 0.213 | 0.725 ± 0.385 | 0.465 ± 0.398 | 0.766 ± 0.205 | 897 |
|  | Minimal + DK + Guide | 0.781 ± 0.328 | 0.853 ± 0.274 | 0.881 ± 0.218 | 0.886 ± 0.235 | 0.844 ± 0.296 | 0.793 ± 0.238 | 0.840 ± 0.218 | 888 |

**Table S4. Average and total extraction costs per figure for each MLLMs.** The computational costs were calculated for the figure-by-figure extraction experiments using Figure + Text (C&M) as the input format and Minimal + Guide + DK as the input prompt. The average token consumption per processing unit was 5,388 tokens. Based on this value, the cost in USD for each model was estimated and summarized in the table. The Average cost represents the mean cost per processing unit, whereas the total cost indicates the cumulative cost of processing 20 papers. The Claude Sonnet models share an identical pricing structure and are therefore presented in a single column.

|  | Gemini 2.5 Pro | Gemini 2.0 Flash | Gemini 1.5 Flash | GPT-5 | GPT-4.1 | GPT-4o | Claude Sonnet series |
| --- | --- | --- | --- | --- | --- | --- | --- |
| Average cost (USD) | 0.00724 | 0.00362 | 0.00087 | 0.08387 | 0.01158 | 0.05493 | 0.01737 |
| Total cost (USD) | 0.40522 | 0.20261 | 0.04863 | 4.69675 | 0.64835 | 3.07588 | 0.97253 |

**Table S5. Comparison of extraction performance between scatter plots and microscopic images.** The average F1-scores under each prompt condition: Minimal, Minimal + DK, Minimal + Guide, and Minimal + DK + Guide when using Gemini 2.5 Pro as the base MLLM and Figure + Text (C&M) as the input format. Abbreviations: temp., temperature.

| Input | MLLM | Prompt | Scatter plots | Microscopic images |
| --- | --- | --- | --- | --- |
| Figure + Text (C&M) | Gemini 2.5 Pro  (Temp. = 0.0) | Minimal | 0.803 | 0.660 |
|  |  | Minimal + DK | 0.791 | 0.707 |
|  |  | Minimal + Guide | 0.838 | 0.776 |
|  |  | Minimal + DK + Guide | 0.848 | 0.849 |
|  | Gemini 2.5 Pro  (Temp. = 0.1) | Minimal + DK + Guide | 0.913 | 0.829 |

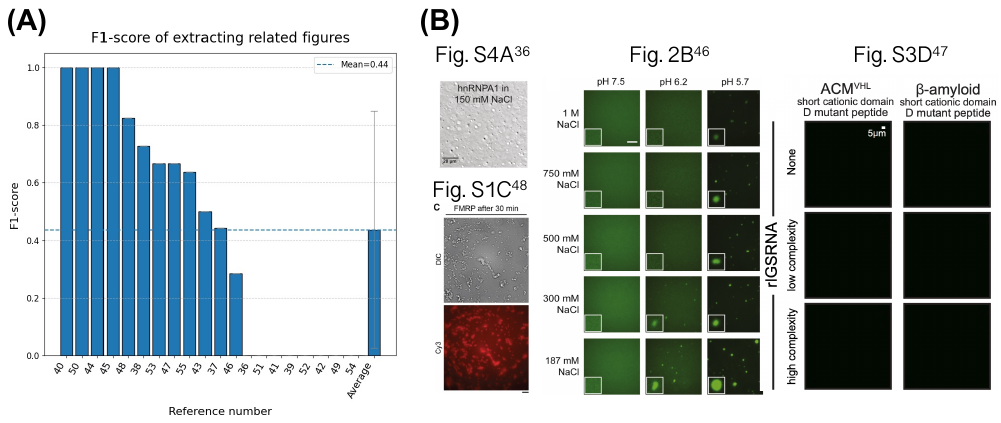

**Figure S1. F1-score and failure examples using automate figure extraction.** To verify whether figure extraction can be automated, we conducted an experiment to extract figure numbers from the PDFs of the 20 evaluation papers included in the evaluation dataset. Similar to single-shot extraction, we input PDFs with added figure number extraction prompts into the MLLM and extracted all relevant figure numbers in bulk. The prompt consisted of two elements: first, requesting an enumeration of all figure numbers within the paper; second, requesting descriptions for each figure, verifying whether they met the extraction criteria, and then compiling a list of all figure numbers satisfying all criteria. Prompt details are documented in the sixth section of the “additional_file2.docx” file. The experiment was conducted using Gemini 2.5 Pro with a temperature parameter set to 0. All the other parameters were maintained at their default values. The extracted results were compared with the target figure numbers in the evaluation dataset. Matches were counted as true positives and the F1 score was calculated. The average F1 score across all the papers served as the final evaluation metric. (A) Average F1-scores for extraction of relevant figure numbers from each paper. The rightmost bar indicates the average across all papers, and the error bars indicate the STD of the average F1-score between each paper. (B) Examples of false positives. Specifically, Fig. S4A^36^ and Fig. 2B^46^ does not contain RNA, Fig. S3D^47^ does not show LLPS, and Fig. S1C^48^ varied the time parameter, which was outside the scope of the extraction. The reference numbers correspond to citations in the main text and Table S1. Therefore, these figures were not included as targets for extraction in this study. Image in Fig. S4A^36^ were reproduced with permission from the paper^36^. Image in Fig. 2B^46^ was reproduced with permission from the paper^46^. Image in Fig. S3D^47^ was reproduced with permission from the paper^47^. Image in Fig. S1C^48^ were reproduced with permission from the paper^48^.
