## Supporting Information (Prompts) for "Assessing the Performance of LLMs in Multimodal Information Extraction for Biological Research: A Case Study on LLPS"

**List of Prompts**

1. **The prompt (a) Minimal used for information extraction.** The placeholder {TARGET_FIGURE_NUMBER} denotes the figure number of the target image for processing. When textual data (e.g., full text, captions, or method descriptions) are included as input, they are appended to the end of this prompt.

Your task is to extract specific experimental conditions used for **Figure {TARGET_FIGURE_NUMBER}** based on the provided information.

=== OUTPUT FORMAT ===

###{fig_num=VALUE, fig_id=VALUE, experimental_result=VALUE, protein_name=VALUE, protein_region=VALUE, protein_modification=VALUE, protein_conc=VALUE, protein_unit=VALUE, rna_name=VALUE, rna_conc=VALUE, rna_unit=VALUE, temperature=VALUE, pH=VALUE, salt_name_conc=SALT_NAME_1: SALT_CONC_1 SALT_UNIT_1; SALT_NAME_2: SALT_CONC_2 SALT_UNIT_2}, {...}, ..., {...}###

=== OUTPUT RULES ===

- Record each set of experimental conditions as a distinct data point ({EXPERIMENT}).

- Do not use a range notation or ambiguous value in numeric fields.

- Only extract values relevant to **Figure {TARGET_FIGURE_NUMBER}**.

- **Do not use '"' or break lines in the final result**.

- **Provide the final result as a single plain sentence**. Do not use JSON or any structured format in other languages.

- The final result **must be one and be enclosed in `###{ ... }###`**. Do not use and `###{` anywhere except for enclosing the final result.

The complete result should exactly follow the output format.

2. **The prompt (b) Minimal + DK used for information extraction.** The placeholder {TARGET_FIGURE_NUMBER} denotes the figure number of the target image for processing. When textual data (e.g., full text, captions, or method descriptions) are included as input, they are appended to the end of this prompt.

Your task is to extract specific experimental conditions used for **Figure {TARGET_FIGURE_NUMBER}** based on the provided information.

=== TARGET CONDITIONS ===

- `fig_num`: Figure number (e.g., 1A).

- `fig_id`: Subfigure label in form "1-a". The x-axis in the figure is represented by numbers, and the y-axis by letters.

- `experimental_result`: Represents the morphology of LLPS-formed condensates, determined based on image or text context. Cannot be "None". Choose one of the following:

- Solute: LLPS did not occur. No objects are visible in the image.

- Liquid: LLPS occurs, forming spherical, smooth-edged liquid-like droplets.

- Gel: LLPS results in interconnected networks or irregular granules.

- Solid: LLPS results in angular, independent granules or aggregates.

- `protein_name`: The **formal and complete protein name** (e.g., FMRP, G3BP1, hnRNPA1).

- Do **not** use abbreviations (e.g., RGG, PLD), tags (e.g., SNAP), or partial sequences (e.g., RGG-3Y, FUS-IDR).

- If a partial domain or tag is mentioned, infer the corresponding full protein name from the context.

- When unclear, prioritize names mentioned in figure titles or methods section over image labels.

- `protein_region`: Sequence region (e.g., Full-length, IDR). Tags are not included.

- `protein_modification`: Modifications (e.g., R10D, acetylated). Tags are not included.

- `protein_conc`: Protein concentration value (numeric only).

- `protein_unit`: Unit of protein concentration (e.g., μM).

- `rna_name`: RNA name.

- `rna_conc`: RNA concentration value (numeric only).

- `rna_unit`: Unit of RNA concentration (e.g., nM).

- `temperature`: Value + unit (e.g., 25°C).

- `pH`: Numeric value. Frequently appears right after the mention of `pH`.

- `salt_name_conc`: List of salts and concentrations, formatted as:

`"NaCl: 150 mM; MgCl2: 5 mM"`

Format and output the extracted results by strictly following the specified **OUTPUT FORMAT** and adhering to all **OUTPUT RULES**:"""+"""

=== OUTPUT FORMAT ===

###{fig_num=VALUE, fig_id=VALUE, experimental_result=VALUE, protein_name=VALUE, protein_region=VALUE, protein_modification=VALUE, protein_conc=VALUE, protein_unit=VALUE, rna_name=VALUE, rna_conc=VALUE, rna_unit=VALUE, temperature=VALUE, pH=VALUE, salt_name_conc=SALT_NAME_1: SALT_CONC_1 SALT_UNIT_1; SALT_NAME_2: SALT_CONC_2 SALT_UNIT_2}, {...}, ..., {...}###

=== OUTPUT RULES ===

- Record each set of experimental conditions as a distinct data point ({EXPERIMENT}).

- Do not use a range notation or ambiguous value in numeric fields."""+f"""

- Only extract values relevant to **Figure {TARGET_FIGURE_NUMBER}**.

- **Do not use '"' or break lines in the final result**.

- **Provide the final result as a single plain sentence**. Do not use JSON or any structured format in other languages.

- The final result **must be one and be enclosed in `###{ ... }###`**. Do not use and `###{` anywhere except for enclosing the final result.

3. **The prompt (c) Minimal + Guide used for information extraction.** The placeholder {TARGET_FIGURE_NUMBER} denotes the figure number of the target image for processing. When textual data (e.g., full text, captions, or method descriptions) are included as input, they are appended to the end of this prompt.

Your task is to extract specific experimental conditions used for **Figure {TARGET_FIGURE_NUMBER}** based on the provided image, caption, and method text.

You must analyze both the **text** (caption and method) and the **image content** (e.g., axis labels, units, legend) to complete this task.

=== INSTRUCTIONS ===

1. Identify all relevant sections related to Figure {TARGET_FIGURE_NUMBER} using all available resources (e.g., image, caption, method text).

2. Based on the figure type, distinguish all individual experiments shown in the image and identify their specific experimental conditions accordingly.

**(A) Scatterplot**

- Each **data point** represents a **distinct experiment**.

- Use **x-axis/y-axis labels and units** to map coordinates to parameters such as `protein_conc`, `rna_conc`, `temperature`, or `pH`.

- Use **legend meaning** (not color or shape alone) to infer `experimental_result` (e.g., Liquid, Solute).

- Prioritize values explicitly mentioned in the image (e.g., axis labels) when text is ambiguous.

**(B) Microscopy Image**

- Even if all images are from the same imaging condition, multiple **distinct experimental conditions** (e.g., different concentrations) may be shown.

- Treat each distinct experimental condition as a separate experiment to extract.

- Determine `experimental_result` from:

- Caption or method description.

- Image morphology (e.g., smooth droplets → Liquid, aggregates → Solid, networks → Gel).

3. Extract all the following fields:

** FIELDS **

- `fig_num`

- `fig_id`

- `experimental_result`

- `protein_name`

- `protein_region`

- `protein_modification`

- `protein_conc`

- `protein_unit`

- `rna_name`

- `rna_conc`

- `rna_unit`

- `temperature`

- `pH`

- `salt_name_conc`

** RULE FOR EXTRACTING FIELDS **

(A) The rule to extract numeric items:

- Each numeric field (`*_conc`, `pH`, `temperature`) must be a **single, specific value**.

- Do **not** use ranges (e.g., "10–20 μM") or vague expressions.

- Instead, extract a representative value (e.g., choose "10 μM").

(B) The following fields may be `None` only if no relevant information is available:

`protein_region`, `protein_modification`, `salt_name_conc`.

- After extracting each value, verify that your interpretation is grounded in the source text or image. If you are uncertain, leave the value as "None".

(C) The following fields must always be extracted and cannot be `None`. Should extract specific value:

`protein_name`, `protein_conc`, `protein_unit`, `rna_name`, `rna_conc`, `rna_unit`, `experimental_result`, `temperature`, `pH`.

- After extracting each value, verify that your interpretation is grounded in the source text or image. If you are uncertain, reassess the context carefully and make your best-supported inference.

"""+"""

4. Format and output the extracted results by strictly following the specified **OUTPUT FORMAT** and adhering to all **OUTPUT RULES**:

=== OUTPUT FORMAT ===

###{protein_name=SPECIFIC VALUE, protein_region=SPECIFIC VALUE, protein_modification=SPECIFIC VALUE, protein_conc=SPECIFIC VALUE, protein_unit=SPECIFIC VALUE, rna_name=SPECIFIC VALUE, rna_conc=SPECIFIC VALUE, rna_unit=SPECIFIC VALUE, experimental_result=SPECIFIC VALUE, temperature=SPECIFIC VALUE, pH=SPECIFIC VALUE, salt_name_conc=SPECIFIC VALUE}, {EXPERIMENT_2}, {EXPERIMENT_3}, ..., {EXPERIMENT_N}###

=== OUTPUT RULES ===

- Record each set of experimental conditions as a distinct data point ({EXPERIMENT}).

- Do not use a range notation or ambiguous value in numeric fields.

- Only extract values relevant to **Figure {TARGET_FIGURE_NUMBER}**.

- **Do not use '"' or break lines in the final result**.

- **Provide the final result as a single plain sentence**. Do not use JSON or any structured format in other languages.

- The final result **must be one and be enclosed in `###{ ... }###`**. Do not use and `###{` anywhere except for enclosing the final result.

Complete the task accurately by following the previous instructions.

Think step by step: locate the data, interpret it, and format the result."""

4. **the prompt (d) Minimal + DK + Guide used for information extraction.** The placeholder {TARGET_FIGURE_NUMBER} denotes the figure number of the target image for processing. When textual data (e.g., full text, captions, or method descriptions) are included as input, they are appended to the end of this prompt.

You are an LLPS domain expert skilled in inferring and validating experimental conditions from visual and textual data.

Your task is to extract specific experimental conditions used for **Figure {TARGET_FIGURE_NUMBER}** based on the provided image, caption, and method text.

You must analyze both the **text** (caption and method) and the **image content** (e.g., axis labels, units, legend) to complete this task.

=== INSTRUCTIONS ===

1. Identify all relevant sections related to Figure {TARGET_FIGURE_NUMBER} using all available resources (e.g., image, caption, method text).

2. Based on the figure type, distinguish all individual experiments shown in the image and identify their specific experimental conditions accordingly.

**(A) Scatterplot**

- Each **data point** represents a **distinct experiment**.

- Use **x-axis/y-axis labels and units** to map coordinates to parameters such as `protein_conc`, `rna_conc`, `temperature`, or `pH`.

- Use **legend meaning** (not color or shape alone) to infer `experimental_result` (e.g., Liquid, Solute).

- Prioritize values explicitly mentioned in the image (e.g., axis labels) when text is ambiguous.

**(B) Microscopy Image**

- Even if all images are from the same imaging condition, multiple **distinct experimental conditions** (e.g., different concentrations) may be shown.

- Treat each distinct experimental condition as a separate experiment to extract.

- Determine `experimental_result` from:

- Caption or method description.

- Image morphology (e.g., smooth droplets → Liquid, aggregates → Solid, networks → Gel).

3. Extract all the following fields:

** FIELDS **

- `fig_num`: Figure number (e.g., 1A).

- `fig_id`: Subfigure label in form "1-a". The x-axis in the figure is represented by numbers, and the y-axis by letters.

- `experimental_result`: Represents the morphology of LLPS-formed condensates, determined based on image or text context. Cannot be "None". Choose one of the following:

- Solute: LLPS did not occur. No objects are visible in the image.

- Liquid: LLPS occurs, forming spherical, smooth-edged liquid-like droplets.

- Gel: LLPS results in interconnected networks or irregular granules.

- Solid: LLPS results in angular, independent granules or aggregates.

- `protein_name`: The **formal and complete protein name** (e.g., FMRP, G3BP1, hnRNPA1).

- Do **not** use abbreviations (e.g., RGG, PLD), tags (e.g., SNAP), or partial sequences (e.g., RGG-3Y, FUS-IDR).

- If a partial domain or tag is mentioned, infer the corresponding full protein name from the context.

- When unclear, prioritize names mentioned in figure titles or methods section over image labels.

- `protein_region`: Sequence region (e.g., Full-length, IDR). Tags are not included.

- `protein_modification`: Modifications (e.g., R10D, acetylated). Tags are not included.

- `protein_conc`: Protein concentration value (numeric only).

- `protein_unit`: Unit of protein concentration (e.g., μM).

- `rna_name`: RNA name.

- `rna_conc`: RNA concentration value (numeric only).

- `rna_unit`: Unit of RNA concentration (e.g., nM).

- `temperature`: Value + unit (e.g., 25°C).

- `pH`: Numeric value. Frequently appears right after the mention of `pH`.

- `salt_name_conc`: List of salts (e.g., NaCl, MgCl2) and concentrations. A salt is a neutral compound formed by the electrostatic attraction between positively charged cations and negatively charged anions. Formatted as:

`"NaCl: 150 mM; MgCl2: 5 mM"`

** RULE FOR EXTRACTING FIELDS **

(A) The rule to extract numeric items:

- Each numeric field (`*_conc`, `pH`, `temperature`) must be a **single, specific value**.

- Do **not** use ranges (e.g., "10–20 μM") or vague expressions.

- Instead, extract a representative value (e.g., choose "10 μM").

(B) The following fields may be `None` only if no relevant information is available:

`protein_region`, `protein_modification`, `salt_name_conc`.

- After extracting each value, verify that your interpretation is grounded in the source text or image. If you are uncertain, leave the value as "None".

(C) The following fields must always be extracted and cannot be `None`. Should extract specific value:

`protein_name`, `protein_conc`, `protein_unit`, `rna_name`, `rna_conc`, `rna_unit`, `experimental_result`, `temperature`, `pH`.

- After extracting each value, verify that your interpretation is grounded in the source text or image. If you are uncertain, reassess the context carefully and make your best-supported inference.

4. Format and output the extracted results by strictly following the specified **OUTPUT FORMAT** and adhering to all **OUTPUT RULES**:

=== OUTPUT FORMAT ===

###{protein_name=SPECIFIC VALUE, protein_region=SPECIFIC VALUE, protein_modification=SPECIFIC VALUE, protein_conc=SPECIFIC VALUE, protein_unit=SPECIFIC VALUE, rna_name=SPECIFIC VALUE, rna_conc=SPECIFIC VALUE, rna_unit=SPECIFIC VALUE, experimental_result=SPECIFIC VALUE, temperature=SPECIFIC VALUE, pH=SPECIFIC VALUE, salt_name_conc=SPECIFIC VALUE}, {EXPERIMENT_2}, {EXPERIMENT_3}, ..., {EXPERIMENT_N}###

=== OUTPUT RULES ===

- Record each set of experimental conditions as a distinct data point ({EXPERIMENT}).

- Do not use a range notation or ambiguous value in numeric fields."""+f"""

- Only extract values relevant to **Figure {TARGET_FIGURE_NUMBER}**."""+"""

- **Do not use '"' or break lines in the final result**.

- **Provide the final result as a single plain sentence**. Do not use JSON or any structured format in other languages.

- The final result **must be one and be enclosed in `###{ ... }###`**. Do not use and `###{` anywhere except for enclosing the final result.

Complete the task accurately by following the previous instructions.

Think step by step: locate the data, interpret it, and format the result.

5. **Prompt for Single-shot extraction.**

"""You are an LLPS domain expert skilled in inferring and validating experimental conditions from visual and textual data.

Your task is to extract specific experimental conditions based on the provided files.

You must analyze both the **text** (e.g., caption, method, result) and the **image content** (e.g., axis labels, units, legend) to complete this task.

=== INSTRUCTIONS ===

1. For each figure panel, determine whether it satisfies **all of the following nine conditions**.

Only extract experimental data from subfigures that answer **YES** to **all** of the following questions:

1. Is the experiment related to the **observation of phase separation (LLPS)**?

- LLPS must be clearly indicated by the image (e.g., droplet morphology) or described in the text (e.g., “condensates formed”, “phase diagram”).

- *Turbidity or absorbance alone is not sufficient. *

2. Is the figure a **microscopy image** or a **scatterplot**?

3. Does the experiment involve **only one type of protein** and **only one type of RNA**?

- Complexes (e.g., fusion proteins, chimeras, RNA duplexes) are considered one type.

- If multiple distinct proteins or RNAs are used simultaneously or cannot be determined, the answer is 'No'.

4. Are the varied experimental parameters limited to the **allowed list**?

- Permitted parameters include:

- `protein_name`, `protein_modification`, `protein_region`, `protein_sequence`, `protein_conc`

- `rna_name`, `rna_sequence`, `rna_conc`

- `temperature`, `pH`, `salt name`, `salt concentration`

- If other variables are changed (e.g., time, DNA, crowding agents), the answer is 'No'.

5. Was the experiment conducted **in vitro**?

- Exclude in vivo or in-cell experiments.

6. Is the subfigure **not simply a summary, visualization, or duplication** of another experiment?

- If the figure is discussed in the main text and the referenced experiment only appears in the supplement, the answer is 'Yes'.

7. Does the experiment aim to observe **static phase behavior** (e.g., presence or absence of LLPS) rather than **dynamic properties**?

8. For **scatterplots**, is information about **phase behavior clearly shown** in the legend or caption (e.g., color, symbol, or shape meaning)?

9. Are the values of all varied parameters (e.g., `protein_conc`, `rna_conc`, `temperature`) **explicitly described** in the image, caption, or method section?

- The figures that only use vague phrases like “high RNA” or “variable protein” are suggested as 'No'.

2. Based on the figure type, distinguish all individual experiments shown in the image and identify their specific experimental conditions accordingly.

**(A) Scatterplot**

- Each **data point** represents a **distinct experiment**.

- Use **x-axis/y-axis labels and units** to map coordinates to parameters such as `protein_conc`, `rna_conc`, `temperature`, or `pH`.

- Use **legend meaning** (not color or shape alone) to infer `experimental_result` (e.g., Liquid, Solute).

- Prioritize values explicitly mentioned in the image (e.g., axis labels) when text is ambiguous.

**(B) Microscopy Image**

- Even if all images are from the same imaging condition, multiple **distinct experimental conditions** (e.g., different concentrations) may be shown.

- Treat each distinct experimental condition as a separate experiment to extract.

- Determine `experimental_result` from:

- Caption or method description.

- Image morphology (e.g., smooth droplets → Liquid, aggregates → Solid, networks → Gel).

3. Extract all the following fields:

** FIELDS **

- `fig_num`: Figure number (e.g., 1A).

- `fig_id`: The location of each panel in the subfigure is represented in the form "1-a". The x-axis in the figure is represented by numbers, and the y-axis by letters.

- `experimental_result`: Represents the morphology of LLPS-formed condensates, determined based on image or text context. Cannot be "None". Choose one of the following:

- Solute: LLPS did not occur. No objects are visible in the image.

- Liquid: LLPS occurs, forming spherical, smooth-edged liquid-like droplets.

- Gel: LLPS results in interconnected networks or irregular granules.

- Solid: LLPS results in angular, independent granules or aggregates.

- `protein_name`: The **formal and complete protein name** (e.g., FMRP, G3BP1, hnRNPA1).

- Do **not** use abbreviations (e.g., RGG, PLD), tags (e.g., SNAP), or partial sequences (e.g., RGG-3Y, FUS-IDR).

- If a partial domain or tag is mentioned, infer the corresponding full protein name from the context.

- When unclear, prioritize names mentioned in figure titles or methods section over image labels.

- `protein_region`: Sequence region (e.g., Full-length, IDR). Tags are not included.

- `protein_modification`: Modifications (e.g., R10D, acetylated). Tags are not included.

- `protein_conc`: Protein concentration value (numeric only).

- `protein_unit`: Unit of protein concentration (e.g., μM).

- `rna_name`: RNA name.

- `rna_conc`: RNA concentration value (numeric only).

- `rna_unit`: Unit of RNA concentration (e.g., nM).

- `temperature`: Value + unit (e.g., 25°C).

- `pH`: Numeric value. Frequently appears right after the mention of `pH`.

- `salt_name_conc`: List of salts (e.g., NaCl, MgCl2) and concentrations. A salt is a neutral compound formed by the electrostatic attraction between positively charged cations and negatively charged anions. Formatted as:

`"NaCl: 150 mM; MgCl2: 5 mM"`

** RULE FOR EXTRACTING FIELDS **

(A) The rule to extract numeric items:

- Each numeric field (`*_conc`, `pH`, `temperature`) must be a **single, specific value**.

- Do **not** use ranges (e.g., "10–20 μM") or vague expressions.

- Instead, extract a representative value (e.g., choose "10 μM").

(B) The following fields may be `None` only if no relevant information is available:

`protein_region`, `protein_modification`, `salt_name_conc`.

- After extracting each value, verify that your interpretation is grounded in the source text or image. If you are uncertain, leave the value as "None".

(C) The following fields must always be extracted and cannot be `None`. Should extract specific value:

`protein_name`, `protein_conc`, `protein_unit`, `rna_name`, `rna_conc`, `rna_unit`, `experimental_result`, `temperature`, `pH`.

- After extracting each value, verify that your interpretation is grounded in the source text or image. If you are uncertain, reassess the context carefully and make your best-supported inference.

"""+"""

4. Format and output the extracted results by strictly following the specified **OUTPUT FORMAT** and adhering to all **OUTPUT RULES**:

=== OUTPUT FORMAT ===

###{fig_num=SPECIFIC VALUE, fig_id=SPECIFIC VALUE, protein_name=SPECIFIC VALUE, protein_region=SPECIFIC VALUE, protein_modification=SPECIFIC VALUE, protein_conc=SPECIFIC VALUE, protein_unit=SPECIFIC VALUE, rna_name=SPECIFIC VALUE, rna_conc=SPECIFIC VALUE, rna_unit=SPECIFIC VALUE, experimental_result=SPECIFIC VALUE, temperature=SPECIFIC VALUE, pH=SPECIFIC VALUE, salt_name_conc=SPECIFIC VALUE}, {EXPERIMENT_2}, {EXPERIMENT_3}, ..., {EXPERIMENT_N}###

=== OUTPUT RULES ===

- Record each set of experimental conditions as a distinct data point ({EXPERIMENT}).

- Do not use a range notation or ambiguous value in numeric fields.

- **Do not use '"' or break lines in the final result**.

- **Provide the final result as a single plain sentence**. Do not use JSON or any structured format in other languages.

- The final result **must be one and be enclosed in `###{ ... }###`**. Do not use and `###{` anywhere except for enclosing the final result.

Complete the task accurately by following the previous instructions.

Think step by step: locate the data, interpret it, and format the result.

Return the reasoning process and the extracted results of all related figures.

6. **Prompt for extracting target figure numbers.**

You are an expert in liquid-liquid phase separation (LLPS), tasked with identifying all figure panels in a scientific paper that meet a strict set of experimental criteria. There is always at least one qualifying figure panel in the paper, so examine each figure carefully and thoroughly.

Follow the steps below in order. Use all available information, including figure images, figure captions, and the methods section.

---

\# INSTRUCTIONS

\## Step 1: List all subfigure numbers

Identify and enumerate all subfigure labels that appear in the paper, such as `1A`, `2B`, `3C`, etc.

---

\## Step 2: Describe each subfigure’s experiment

For each subfigure listed in Step 1, briefly describe the experimental content based on the figure, caption, and method text. Address the following nine aspects for each subfigure:

1. Is the experiment related to the **observation of phase separation (LLPS)**?

- LLPS must be clearly indicated either by the image (e.g., droplet morphology) or the text (e.g., condensate formation, phase diagram).

- *Turbidity or absorbance measurement alone is not sufficient.*

2. Is the figure a **microscopy image** or a **scatterplot**?

- *Line plots such as time-course or absorbance curves are not considered scatterplots and must be excluded.*

3. Does the experiment involve **only one type of protein** and **one type of RNA**?

- Complexes (e.g., fusion proteins, chimeras, RNA duplexes) count as one type.

- If multiple components are used simultaneously or cannot be identified, exclude the subfigure.

4. Are the experimental parameters varied among the **allowed list** only?

- Permitted items include:

- `protein_name`, `protein_modification`, `protein_region`, `protein_sequence`, `protein_conc`

- `rna_name`, `rna_sequence`, `rna_conc`

- `temperature`, `pH of buffer`, `salt name`, `salt concentration`

- If the subfigure varies unlisted parameters (e.g., time, DNA, crowding agents), it must be excluded.

5. Was the experiment conducted **in vitro**?

- In vivo or in-cell experiments must be excluded.

6. Is the subfigure **not simply a summary, visualization, or duplication** of another experiment?

- However, if it appears in the main text and the referenced experiment is only in the supplementary figures, include it.

7. Is the experiment designed to observe **static phase behavior at a specific condition or timepoint**, rather than **dynamic properties** (e.g., viscosity, elasticity, FRAP, microrheology)?

8. For **scatterplots**, is information about phase behavior (e.g., presence/absence of droplets) **clearly indicated** in the legend or caption using color, shape, or symbol?

9. Are the values of `protein_conc`, `rna_conc`, and any other varied parameters **explicitly described** in the figure, caption, or method section?

- Do not include figures with vague or qualitative statements (e.g., “high RNA” or “variable protein”).

---

\## Step 3: Evaluate each subfigure against all criteria

For each subfigure, answer the following **nine Yes/No questions**, details corresponding directly to the nine items from Step 2:

1. Is there clear evidence this is about **LLPS**?

2. Is the **figure type microscopy or scatterplot**?

3. Are **exactly one protein and one RNA** used in each individual experiment? However, If the majority of experiments in the subfigure involve one type of protein and one type of RNA, and only a minority involve either protein only or RNA only, you may classify it as yes.

4. Are the varied parameters limited to the **allowed list**?

5. Was the experiment conducted **in vitro**?

6. Is this subfigure **not** a summary, visualization or duplication of other figures? If this subfigure is shown in the main text and the referenced experiment is only in the supplementary figures, consider as **yes**.

7. Is this experiment focused on static phase observation, **not dynamic measurements** like FRAP or microrheology?

8. If this is a **scatterplot**, are phase-related outcomes clearly described in the legend or caption?

9. Are both protein concentration and RNA concentration in all experiments of this figure can be found in the text or image?

> If **any answer is No or unclear**, exclude the subfigure.

---

\## Step 4: Output valid subfigures using the specified format

- If one or more subfigures satisfy **all nine conditions**, output:

`+++1A, 2B, 3C+++`

---

\# NOTES

- Use all available sources (figures, captions, methods).

- Do **not** guess. Select only if criteria are explicitly satisfied.

- Be strict. Do not include subfigures with partial or uncertain compliance.

- You must complete **Steps 1–4** and provide a final output.

- **Only the final result in Step 4 should be enclosed with `+++...+++`. **
